# Learning the payoffs and costs of actions

**DOI:** 10.1101/346114

**Authors:** Moritz Möller, Rafal Bogacz

## Abstract

A set of sub-cortical nuclei called basal ganglia is critical for learning the values of actions. The basal ganglia include two pathways, which have been associated with approach and avoid behavior respectively, and are differentially modulated by dopamine projections from the midbrain. According to the influential opponent actor learning model, these pathways represent learned estimates of the positive and negative consequences (payoffs and costs) of actions. The level of dopamine release controls to what extent payoffs and costs enter the overall evaluation of actions. How the knowledge about payoff and cost is acquired is still an open question, even though many theories describe learning from feedback in the basal ganglia. We examine whether a set of plasticity rules proposed to model reinforcement learning in the pathways of the basal ganglia is suitable to extract payoffs and costs from a reward prediction error signal. First, we determine the result of such learning, both analytically and via simulations, for different reward schedules that feature payoffs and costs. Then, we combine the plasticity rules with a decision rule to examine the emerging effect of dopaminergic modulation on the willingness to work for reward. We find that the plasticity rules are suitable to infer the mean payoffs and costs of actions, if those occur at different moments in time. Successful learning requires differential effects of positive and negative reward prediction errors on the two pathways, and a weak decay of synaptic weights over trials. We also confirm that dopaminergic modulation produces effects on the willingness to work for reward similar to those observed in classical experiments.

**Author summary:** The basal ganglia are structures underneath the surface of the vertebrate brain, associated with error driven learning. Much is known about the anatomical and biological features of the basal ganglia; scientists now try to understand the algorithms implemented by these structures. Numerous models aspire to capture the learning functionality, but many of them only cover some specific aspect of the algorithm. Instead of further adding to that pool of partial models, we unify two existing ones - one which captures what the basal ganglia learns, and one that describes the learning mechanism itself. The first model suggests that the basal ganglia keeps track of both positive and negative consequences of frequent opportunities, and weighs these by the motivational state in decisions. It explains how payoff and cost are represented, but not how those representations arise. The other model consists of biologically plausible plasticity rules, which describe how learning takes place, but not how the brain makes use of what is learned. We show that the two theories are compatible. Together, they form a model of learning and decision making that integrates the motivational state as well as the learned payoffs and costs of opportunities.

## Introduction

What guides rational behavior in a complex environment? Certainly, knowledge of the typical payoffs and costs of available actions is critical for successful action selection. If those payoffs and costs are represented separately in the animal’s brain, they can be weighted depending of animal’s motivational state. For example, consider the action ‘harvesting a fruit from a tree’. It has a payoff connected with the nutrients in the fruit, but also costs related to the effort and risks associated with climbing a tree. The nutrients in the fruit are only valuable for the animal if its hungry. So, when it is hungry, the payoffs should be weighted more than the costs, to ensure that the animal searches for food. By contrast when the animal is not hungry at all, the costs should be weighted more, to make sure that it does not climb the tree without necessity.

In all vertebrates, an important role in this process of action evaluation and selection is played by a set of subcortical structures called the basal ganglia [1]. The basal ganglia is organized into two main pathways shown schematically in green and red in Fig 1. The Go or direct pathway is related to the initiation of movements, while the No-Go or indirect pathway is related to the inhibition of movements [2]. These two pathways include two separate populations of striatal neurons expressing different dopaminergic receptors [3]. The striatal Go neurons express D1 receptors and are excited by dopamine, while the striatal No-Go neurons express D2 receptors and are inhibited by dopamine [4]. Thus dopamine changes the balance between the two pathways and promotes action initiation over inhibition.

**Fig 1.**
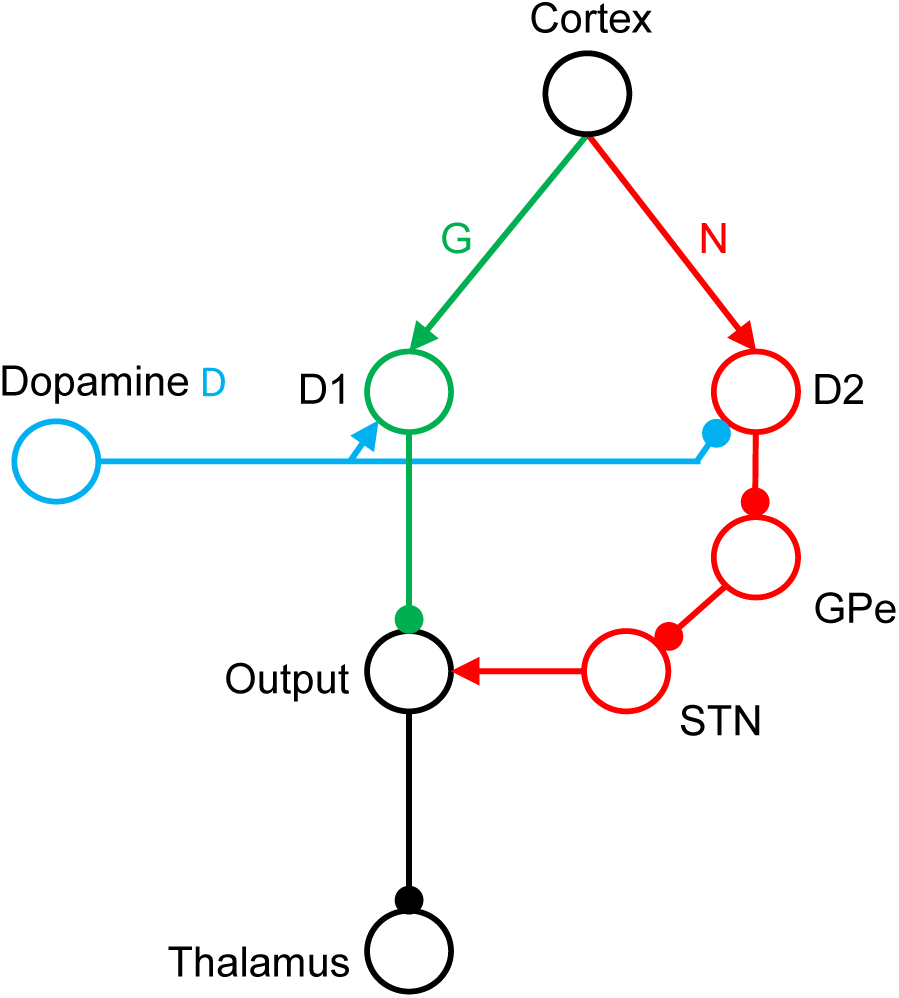
The organization of the basal ganglia. Circles denote neural populations in the areas indicated by labels next to them, where D1 and D2 corresponds to striatal neurons expressing D1 and D2 receptors respectively, STN stands for the subthalamic nucleus, GPe for the external segment of globus pallidus, and Output for the output nuclei of the basal ganglia, i.e. internal segment of globus pallidus and substantia nigra pars reticulata. Arrows and lines ending with circles denote excitatory and inhibitory connections respectively.

The competition between Go and No-Go pathways during action selection and the role of dopaminergic modulation are subject of many interpretations and models, e.g. [5–7]. In particular, the Opponent Actor Learning (OpAL) hypothesis suggests that the Go and No-Go neurons encode the positive and negative consequences of actions respectively [8]. As the dopaminergic neurons modulate the Go and No-Go neurons in opposite ways, dopamine controls the extent to which positive and negative consequences affect the activity in the thalamus, through the output of the basal ganglia [8]. For example, when motivation is high, the dopaminergic neurons will excite the Go neurons and inhibit the No-Go neurons. Consequently, the payoffs will be weighted stronger than the costs. By contrast, when the motivation is low, the Go neurons will not be excited, but the No-Go neurons will be released from inhibition, such that the costs are weighted stronger.

Much research has also focused on how the synapses of Go and No-Go neurons are modified by experience. Systematic investigation revealed that bursts of activity of dopaminergic neurons encode reward prediction errors, which measure the difference between reward obtained and expected [9, 10]. Such dopaminergic activity produces distinct changes in the synaptic weights of Go and No-Go neurons [11]. Several computational models have attempted to describe the learning process of the synapses of Go and No-Go neurons [12–15]. Among these models, the OpAL model provided simple and analytically tractable rules describing the changes in weights of Go and No-Go neurons as a function of reward prediction errors [8].

However, no-one so far examined how the basal ganglia might estimate payoff and cost if they are both associated with the same action. The novel contribution of this work is to demonstrate that a set of recently proposed learning rules [16] allows the Go and No-Go neurons to estimate both payoffs and costs associated with single actions. We thus merge the interpretation of the striatal pathways of Collins and Frank with the striatal learning rules of Mikhael and Bogacz, to ultimately obtain a consistent theory of learning the payoffs and costs of actions.

According to the experimental and modelling work mentioned above, dopaminergic activity encodes both information about motivational state and the reward prediction error. However, if the dopaminergic neurons carried both signals, the striatal neurons would need a way to decode each signal and react appropriately, i.e. change their activity according to the motivation signal, and change the synaptic weights according to the prediction error. The prominent suggestion that motivation might be encoded in the average or tonic dopamine level, and reward prediction errors in the burst or phasic activity [17] has recently been questioned; it seems to be contradicted by the observation of fast-changing dopaminergic activity that encodes motivation [18–20]. Nevertheless, the motivation and teaching signals could both be provided by other means. For example, the activity of striatal cholinergic neurons may inform what the dopaminergic neurons encode at the moment. Such a role of acetylcholine is consistent with the observation that the cholinergic interneurons pause when feedback is provided [21], with the models of intreacellular pathways suggesting that the reduced concentration of acetylcholine is necessary for the striatal plasticity [22], and with other data reviewed elsewhere [20]. In this paper we assume that striatal neurons can read out both motivational and teaching signals encoded by dopaminergic neurons, and we leave the details of the mechanisms by which they can be distinguished to future work.

## Results

Following the OpAL model [8], we assume that the positive consequences of actions are encoded by synaptic weights within the Go pathway. More precisely, we claim that the typical payoff of a particular action in a particular situation is encoded in the strength of the connections from the cortical neurons selective for the situation to the striatal Go neurons selective for the action. We denote these weights by *G* (see Fig 1), and propose that after learning, the weights *G* represent the mean payoff for an action. Mathematically, the collective strength of the weights *G* corresponds to a single, non-negative number. The negative consequences, on the other hand, are encoded in the synaptic connections of striatal No-Go neurons. We denote their weights by *N*, and propose that after learning, they represent the mean cost of an action. Just as with *G*, we mathematically represent the collective strength of the weights *N* by a single, non-negative number.

To learn the positive and negative consequences of actions respectively, the striatal neurons can take advantage of the fact that these consequences typically occur in different moments in time. Let us consider a situation in which an animal performs an action that involves an effort in order to obtain a reward: Fig 2a sketches a task in which a rat is given the opportunity to press a lever in order to obtain a food pellet. Due to the effort, the instantaneous reinforcement during the course of this action is negative at first, while pressing the lever. Then, it turns positive at the time the payoff is received. Fig 2b sketches the resulting changes in the synaptic weights. The leftmost display shows the initial weights. While making an effort to perform an action, the reward prediction error is negative. Similarly as in previous models [8, 12], we assume that the negative prediction error results in a strengthening of *N* (compare the red arrows in the middle and the left displays in Fig 2b). This allows the weights *N* to encode negative consequences. Later, reception of the payoff causes a positive prediction error, which strengthens *G*. This leads the weights *G* to encode the positive consequences. So if an experience involves both positive and negative consequences, both weights are increased during the experience (compare the right and the left displays in Fig 2b).

**Fig 2.**
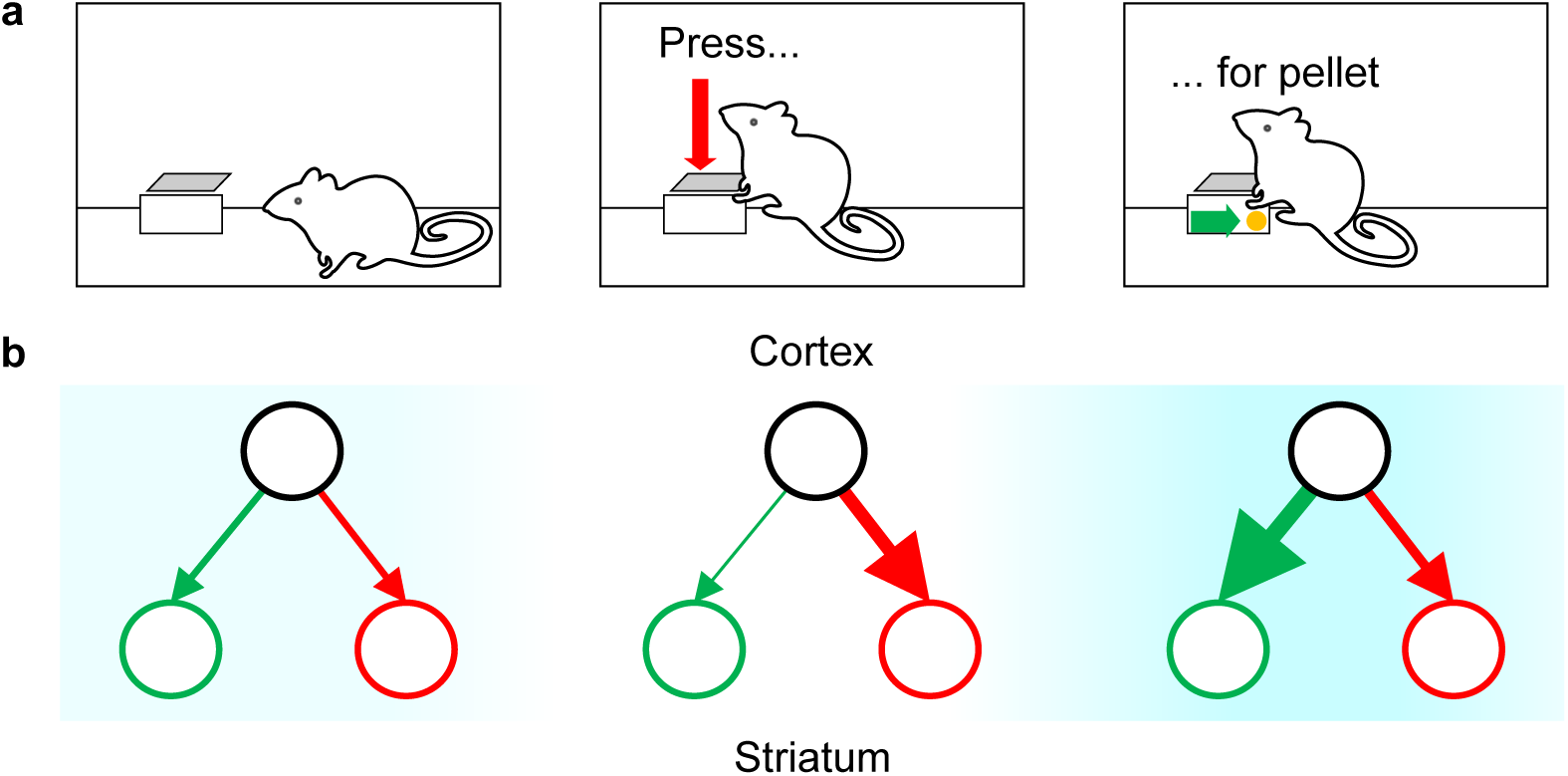
Qualitative description of learning payoffs and costs. (a) Operant conditioning chamber setup: a rat obtains a food pellet by pressing a lever. (b) Diagrams of changes in the weights *G* and *N* associated with lever pressing at each stage of the experience presented in panel (a). In all diagrams, the black circles represent the cortical neurons selective for the state (being in the operant box), and the green and red circles represent the Go and No-Go populations of striatal neurons, respectively, selective for the action (pressing lever). The thickness of the arrows linking the circles represents the connection strength between the respective neuron populations. The blue shading in the background indicates the strength of the immediate reinforcement, with color intensity proportional to the magnitude of reward.

To mathematically implement these ideas, we need to model the weighs of the Go pathway *G*, the weighs of the No-Go pathway *N*, and the prediction error. The reward prediction error, which we denote by *δ*, quantifies the difference between the expected reward and the received reward *r*. If *r* is negative, we shall speak of cost, and when *r* is positive, we shall speak of payoff. Payoff creates positive reinforcement, and thus attraction, whilst cost creates negative reinforcement, and thus avoidance. The expected reward, on the other hand, directly corresponds to the expected payoffs and costs, which - according to our theory - are represented by the synaptic weights *G* and *N*. We take the expected reward to be the average over the expected payoff and the expected cost. All together, we model the reward prediction error as

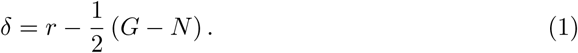

Equipped with the quantities *δ*, *G* and *N*, we can formulate our theory of learning payoff and cost. To do so, we simply describe how the collective connection strengths *G* and *N* change when a prediction error *δ* is received; we use Δ*G* and Δ*N* to denote the changes in connection strengths. Note that any update only applies if the resulting weights are still positive - if an update would render any one weight negative, that weight is set to zero instead. In all other cases, we follow Mikhael and Bogacz [16] in prescribing

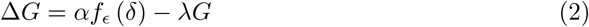

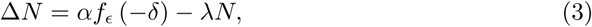

where *α* is the learning rate, *ϵ* is the slope parameter and *λ* the decay rate. The slope parameter *ϵ* controls the strength of the nonlinearity exhibited by the function *f_ϵ_*, which we introduce in Fig 3d - e. The contribution of this article is to point out that if *G* and *N* change according to these rules, they will eventually represent payoff and cost.

**Fig 3.**
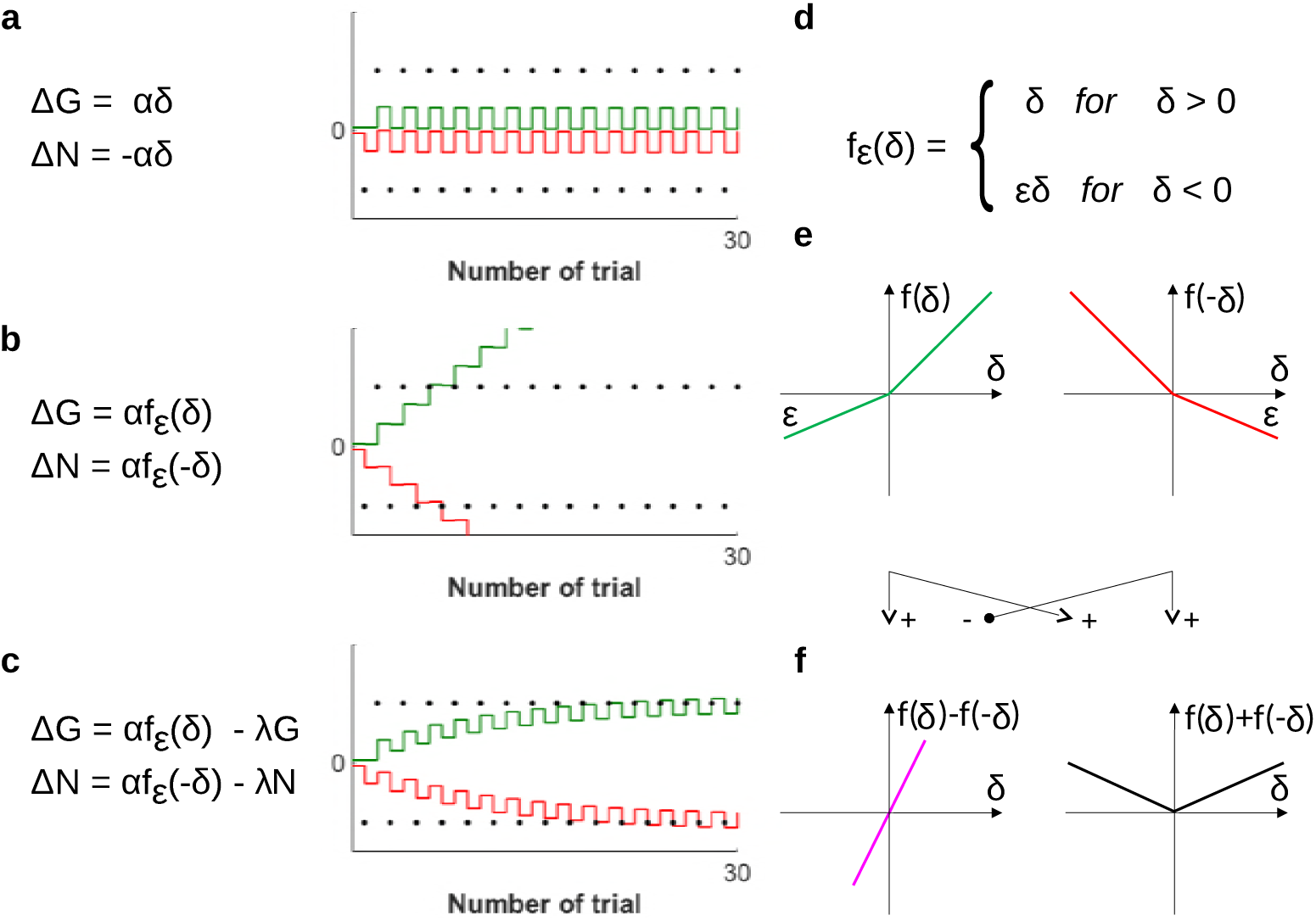
The incremental construction of the learning rules. (a) - (c) The different stages in the construction of the learning rules. All panels feature a mathematical formulation of the rules at the given stage, and a simulation of these rules. The rewards in those simulations, indicated by black dots, alternate between a fixed payoff of magnitude 20 and a fixed cost of −20. The Go weights *G* are depicted in green, the negative No-Go weights −*N* are depicted in red. The parameters used in the simulations were *α* = 0.300, *ϵ* = 0.443 and *λ* = 0.093. (d) - (f) Definition, visualization and properties of the nonlinear function *f_ϵ_*.

There is an intuition for each term in the rules 2 and 3. These intuitions are most easily gained by constructing the rules from scratch; therefore we will now retrace the three steps of that construction. Several models of learning in Go and No-Go neurons assume that the effect of the prediction error on *G* is opposite to its effect on *N* [7, 8]. We thus start by proposing that Δ*G* and Δ*N* might simply be proportional to the prediction error and its negative, respectively. To see whether this proposal works, we formulate it mathematically and simulate the learning of an alternating sequence of costs −*n* and payoffs *p*. Fig 3a shows both the mathematical formulation and the simulation. There is a problem: the strengthening of *N* due to negative prediction error, caused by the cost, is always immediately reversed by the following positive prediction error caused by the payoff. The same is true for the changes in *G*. As illustrated by the simulation, there is no net effect of learning.

To overcome this problem, we proceed by damping the impact of negative prediction errors (which are usually caused by costs) on *G*, and the impact of positive prediction errors on *N*. This is logical, since costs should not alter the estimate *G* of the payoffs, and vice versa. Such damping can be achieved by replacing the simple proportionality to *δ* in the first proposal by a nonlinear dependence, mediated by the functions depicted in Fig 3e. We update our mathematical formulation accordingly, and again simulate the effects of the previously used reward sequence - both these steps are illustrated in Fig 3b. The simulation shows that, while producing the appropriate tendencies, these rules cause unconstrained, ongoing strengthening of both connections. Such dynamics are neither biologically plausible, nor useful to infer the actual payoff and cost.

Finally, to stop unconstrained strengthening and stabilize the weighs, we balance growth with decay. Adding decay terms to the mathematical formulation of the rules yields their final form 2 and 3. The simulation in Fig 3c suggests that the construction was successful: the final version of the rules allows the weights to converge to *p* and *n* respectively.

### Mathematical analysis

After providing an intuitive understanding of the learning rules and their mathematical formulation, we proceed to a more rigorous analytical treatment. We saw the potential of Mikhael and Bogacz’ [16] rules to learn payoffs and costs. Appropriate choice of parameters is key to unlock that potential, and we shall now investigate how that choice must be made. In particular, we will derive certain relations between parameters that must be satisfied for payoff and cost to be learned.

Originally, the rules 2 and 3 were meant to describe learning of reward statistics. Mikhael and Bogacz [16] showed that after learning, particular combinations of *G* and *N* will encode the mean 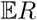 and the mean spread 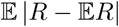 of the received rewards. For further reference, we denote these important statistics by 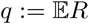 and 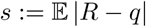. How are the mean and the mean spread of received rewards related to payoff and cost? Consider the reward statistics of an action that reliably requires effort to produce a payoff. Repeat that action multiple times, and record all received rewards, 1 the costs as well as the payoffs. Finally, analyze how all these received rewards are distributed. If effort was required to earn the payoff, the distribution of rewards will turn out bimodal, as schematically shown in Fig 4. It features two peaks, one centered around the mean payoff *p*, and one centered around the mean cost −*n*, respectively. Fig 4 also shows the mean *q* and the mean spread *s* of that distribution. We observe that payoffs and costs are both exactly one mean spread *s* away from the center *q* of the distribution - the payoff above, and the cost below. This implies that there is, at least in this representative case, a strong connection between payoffs and costs and the reward statistics:

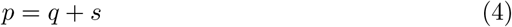

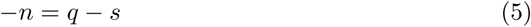

**Fig 4.**
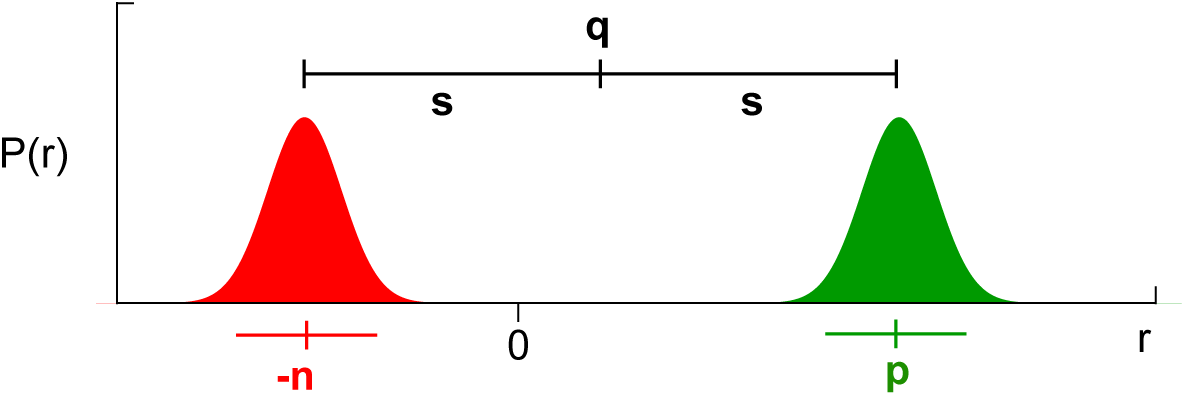
The relation of reward statistics to payoff and cost. The graph shows a representative reward distribution over the magnitude *r* of all received rewards. The parts of the distribution that indicate negative rewards (costs) are colored red, while the parts that indicate positive rewards (payoffs) are colored green. The mean *q* and the mean spread *s* are indicated above the distribution, the mean cost −*n* and the mean payoff *p* are indicated below the distribution.

This connection allows us to set up conditions for the result of learning: if *G* and *N* are to represent payoff and cost, they must approach *q* + *s* and −*q* + *s* respectively. Equivalently, we can ask for 1/2 (*G* − *N*) and 1/2 (*G* + *N*) to approach *q* and *s* in the course of learning.

After revealing the link between reward statistics and payoff and cost, we are ready to derive the relations necessary to learn the latter. To that end, we first determine the connection strengths *G* and *N* that result from training on stochastic rewards. Such uncertain rewards are sampled at random from a fixed distribution. Then, we implement the newly identified conditions, demanding for 1/2 (*G* − *N*) to approximate *q* and 1/2 (*G* + *N*) to approximate *s* after training is finished. From these conditions, we will be able to derive the desired parameter relations.

Working through these steps is simpler after changing variables from *G* and *N* to *Q*:= 1/2 (*G* − *N*) and *S*:= 1/2 (*G* + *N*) right away. We saw that the new variables *Q* and *S* have a clear computational interpretation: if learning goes as planned, *Q* and *S* track the mean *q* and the mean spread *s* of the experienced reward. To determine how *Q* and *S* change due to prediction errors *δ*, we simply add and subtract the update rules 2 and 3. Certain convenient properties of the nonlinear functions *f_ϵ_* help to further simplify the resulting equations: Fig 3f shows that subtracting and adding functions depicted in Fig 3e give functions proportional to identity and absolute value, respectively.^1^ Exploiting these properties, we obtain

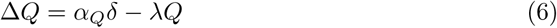

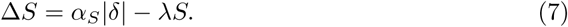

Here, for brevity of notation, we introduced the effective learning rates *α_Q_* = *α* (1 + *ϵ*) /2 and *α_S_* = *α* (1 − *ϵ*)/2. Note that the changes of *Q* and *S* are proportional either to the prediction error itself or to its absolute value, in contrast to the changes of *G* and *N*.

Now, let us determine the strengths of the weights *G* and *N*, or equivalently of the variables *Q* and *S*, after many encounters with an action. When learning the rewards of a previously unknown action, *Q* and *S* typically change a lot during the first trials. These changes then get smaller and smaller as more experience is integrated - the learning curve plateaus. After enough trials, *Q* and *S* stop changing systematically, and start to merely fluctuate about some constant values, which we denote by *Q*^∗^ and *S*^∗^ and refer to as equilibrium points. In mathematical terms, directed learning stops when we may expect *Q* and *S* to remain unchanged by another trial, i.e. when 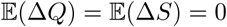. If that stage is reached, the equilibrium points can be inferred by computing the mean value of the fluctuating variables: 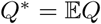 and 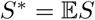. With these identities and the learning rules 6 and 7, we can determine the equilibrium points *Q*^∗^ and *S*^∗^:

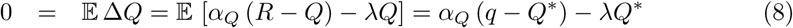

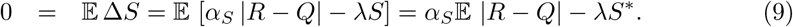

To solve these equations, we shall make the additional assumption that the fluctuations of *Q* about *Q*^∗^ are small. This assumption is justified whenever *α* is sufficiently small, and allows us to approximate 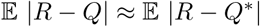. Collecting all those intermediate results, we may solve 8 and 9 for the equilibrium points. The solutions read

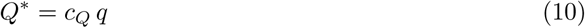

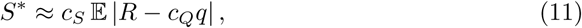

with *c_Q_* = *a_Q_/* (*α_Q_* + *λ*) and 1*/c_S_* = *α_S_ /λ*. Those are the approximate values of *Q* and *S* after learning.

Next, we need to implement the conditions we inferred from Fig 4. Thanks to our choice of variables, this simply amounts to requiring that *Q* converge to the mean reward *q*, and *S* to the mean spread *s*, i.e. requiring *Q*^∗^ = *q* and *S*^∗^ = *s*. Inserting the approximate values 10 and 11 produced by the learning rules, we obtain

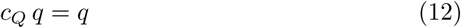

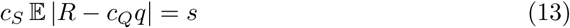

These equations are central to this publication. Their left hand side represents the result of learning according to Mikhael and Bogacz’ [16] rules. Their right hand side specifies what needs to be learned if *G* and *N* really represented payoffs and costs, as Collins and Frank hypothesized [8]. Equating the left and the right hand side amounts to merging both theories. It allows us to determine how the parameters would be related if both theories were exactly true: for 12 and 13 to hold, *α*, *β* and *ϵ* must take values such that *c_Q_* = 1 and *c_S_* = 1.

This result evokes several questions: Is it at all possible to satisfy the derived conditions? What do the conditions mean with respect to the parameters *α*, *λ* and *ϵ*? And finally, is there a practical way to determine sets of parameters *α*, *λ* and *ϵ* which - at least approximately - satisfy the conditions? We discuss each of these questions in the following paragraphs.

Firstly, is it possible to satisfy *c_Q_* = 1 and *c_S_* = 1 exactly? Examining the definition *c_Q_* = *α_Q_/*(*α_Q_* + *λ*) quickly reveals that letting *c_Q_* → 1 would amount to letting *λ* → 0^2^. Now, we derived above that after learning, *S* will fluctuate about its equilibrium point 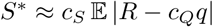 with *c_S_* = *α_S_ /λ*. In order to keep the equilibrium point *S*^∗^ finite as *λ* → 0, we would therefore be forced to have *α_S_* → 0 also. This, though, would pose a real problem: *α_S_* is the effective learning rate for *S* - having it vanish would imply stopping learning in *S* all together. We must conclude that strict satisfaction of the constraints *c_Q_* = 1 and *c_S_* = 1 is not compatible with non-vanishing learning rates that lead to a finite equilibrium. Specifically, *c_Q_* = 1 can only ever hold approximately if the spread *s* is to be learned in finite time. Nevertheless, no such problem arises when *c_S_* is set to 1 exactly.

Now, what do the constraints *c_Q_* ≈ 1 and *c_S_* = 1 mean in terms of the parameters *α*, *λ* and *ϵ* ? In the previous paragraph, we saw that *c_Q_* ≈ 1 is equivalent to *λ/α_Q_* ≈ 0. Since both *λ* (a decay constant) and *α_Q_* (an effective learning rate) are inherently positive, we may rewrite this as *λ/α_Q_* ≪ 1. Inserting the definition *α_Q_* = *α* (1 − *ϵ*) /2 immediately yields

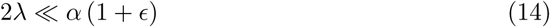

The other condition, *c_S_* = 1, is easily translated analogously. We need only use the definitions *c_S_* = *α_S_ /λ* and *α_S_* = *α* (1 − *ϵ*) /2 to obtain

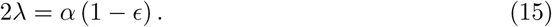

Equations 14 and 15 constitute the exact relations between the parameters *α*, *γ* and *ϵ* that need to hold for payoffs and costs to be estimated accurately. They cannot be further simplified, but we may use them to gain some more insight into the required magnitudes of the individual parameters: by substituting 2*λ* according to Eq 15 on the right hand side of Eq 14, one quickly reaches the conclusion that *ϵ* ≈ 1. Reinserting this into Eq 14 yields *λ* ≪ *α*. In conclusion, we found that it is necessary (though not sufficient) for accurate learning of payoffs and costs to maintain a small, but non vanishing nonlinearity *ϵ* in the transmission of the prediction error signal, as well as a non vanishing decay rate *λ*, which is much smaller than the learning rate *α*.

Finally, how can such parameters *α*, *λ* and *ϵ* practically be determined? To implement the conditions *c_Q_* ≈ 1 and *c_S_* = 1, one can for instance express *λ* and *ϵ* in terms of *α*, *c_Q_* and *c_S_*. It is straight forward to invert the definitions of *c_Q_* and *c_S_* in order to yield *ϵ* = (1 − *c_S_* (1*/c_Q_* − 1))/(1 + *c_S_* (1*/c_Q_* − 1)) and *λ* = *α*(1 − *ϵ*)/(2*c_S_*). Then, one chooses *α* freely at one’s convenience, and *c_Q_* and *c_S_* close (or, in case of *c_S_*, equal) 2 to one. Importantly, *c_Q_* must be chosen smaller then one to result in a positive *λ*. From these choices, one finally obtains *ϵ* and *λ* to work with the chosen *α*. Our simulations suggest that even values such as *c_Q_* = 0.7 and *c_S_* = 0.9, in combination with a learning rate of, say *α* = 0.3, are close enough to one to allow reasonably accurate estimations of payoff and cost. This can be seen in Fig 3: the simulations shown in there used those exact settings, which equivalently means that *ϵ* = 0.443 and *λ* = 0.093.

In summary, we used a statistical argument - the connection between payoffs and costs and the reward statistics - to determine conditions under which payoffs and costs can be learned with the update rules 2 and 3.

### Deterministic reward sequences

In the preceding section, we derived relations that are necessary for successful learning of payoff and cost. If rewards are awarded stochastically, those relations are also sufficient for successful learning. But what happens to the weighs *G* and *N* if the received rewards follow a strong pattern? Assume, for instance, that an action reliably yields a fixed cost −*n* followed by a fixed payoff *p*. Under which additional conditions do *G* and *N* then still reflect the magnitudes of payoff and cost after learning?

To answer that question, we must again determine the connection strengths that result from experiencing the action time and again. Now, we do not have to rely on a probabilistic treatment - when the pattern of the rewards is fully known, it is possible to determine the evolution of *G* and *N* exactly. As in the previous section, we will concentrate on the result of learning rather than on its dynamics. Here, this amounts to determine the fixed points of the learning rules. These fixed points are simply those values of *G* and *N* (or equivalently of the alternative variables *Q* and *S* we defined above) that are invariant under the updates caused by the action. We denote the fixed points by *G*^∗^ and *N*^∗^, or *Q*^∗^ and *S*^∗^. During learning, the variables converge to their respective fixed points, and cease to change notably once they arrive in their vicinity.

First, we focus on determining the fixed point of *Q*. Note that each encounter with the action yields two updates of *Q*: one due to the cost and one due to the payoff. Mathematically, we can formulate this as

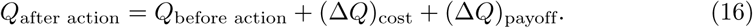

To find *Q*^∗^, demand that these successive updates have no net effect on *Q*: If *Q*_after action_ equals *Q*_before action_, then *Q*_before action_ can rightfully be called fixed point. If this is so, the two updates must have canceled each other:

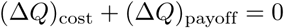

This condition, in combination with the update rules 2 and 3, allows to determine *Q*^∗^ in terms of *p*, *n* and the parameters *α*, *ϵ* and *λ*. One substitutes (Δ*Q*)_cost_ and (Δ*Q*)_payoff_ according to the update rules (note that the *Q* entering the second update had already been changed by the first update), and then solves the equation for *Q*. A straight forward calculation yields

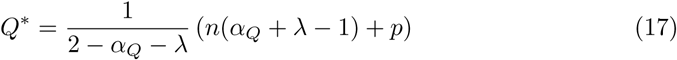

where *α_Q_* = *α* (1 + *ϵ*) /2. Now, recall that the definition of *Q* in terms of *G* and *N* is *Q* = 1/2(*G* − *N*), and that true payoffs and costs of in this model are *p* and *n*. If *G* and *N* represented the true payoffs and costs after learning, it must be true that *G*^∗^ ≈ *p* and *N*^∗^ ≈ *n*, and thereby

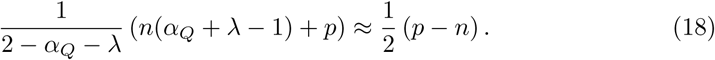

Just as equations 12 and 13, this equation is an interface between the results of Mikhael and Bogacz’ [16] update rules on the left hand side, and Collins and Frank’s hypothesis [8] on the right hand side. For both sides to agree, we must have

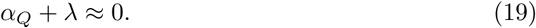

This is a novel condition for learning the correct magnitudes of payoffs and costs from a deterministic reward pattern. The definition of *α_Q_* and the previously derived conditions 14 and 15 may be used to transform this novel condition into the simpler form *α* ≪ 1.

Next, we repeat the same analysis for *S*. Since we search for additional conditions on the parameters, we are free to use the original conditions 14 and 15 to simplify our calculations. The only complication we encounter is the appearance of *Q* in the update rules of *S*, which we resolve by substituting *Q* with *Q*^∗^, acknowledging that the fixed points of *S* and *Q* depend on each other. We arrive at

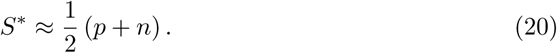

Again, using the definition *S* = 1/2(*G* + *N*) allows to compare the result of learning with the the strengths required to represent payoffs and costs. We immediately find that *G*^∗^ ≈ *p* and *N*^∗^ ≈ *n* already hold. Thus, 19 is the only additional condition for successful learning of payoff and cost from rewards that follow a strong pattern.

From the results presented in this section, we conclude that the learning rules 2 and 3 facilitate learning of the magnitudes of fixed payoffs and costs that occur reliably one after the other. However, we also saw that this is only true if 19 holds in addition to the conditions that we derived in the previous section.

### Summary of analytic results

The analysis above revealed the conditions under which the striatal plasticity rules 2 and 3, put forward by Mikhael and Bogacz [16], could serve the hypothetical function of the striatum proposed by Collins and Frank [8]: to represent the magnitudes of the payoffs and costs of actions. We identified the conditions in two different paradigms: first, we investigated learning from purely stochastic rewards sampled from a fixed distribution. Then, we considered a deterministic pattern of rewards. We obtained two key results:

- Consider a reward distribution - obtained from multiple encounters with an action - that is shaped by payoffs and cost, as the one shown in Fig 4. If trained on rewards sampled from that distribution, the plasticity rules 2 and 3 will enable learning of the mean payoffs and costs if

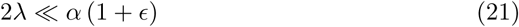

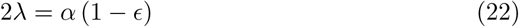

hold. These conditions imply, but do not follow from, a non-vanishing but small nonlinearity in the transmission of the prediction error, and a non-vanishing but small^3^ decay of the connection weights.
- If trained on a pattern of rewards that alternates between payoffs of magnitude *p* and costs of magnitude *n*, the plasticity rules 2 and 3 will capture the those exact payoffs and costs if, in addition to 21 and 22,

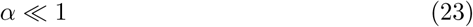

holds. In words, unbiased learning of payoffs and costs in deterministic scenarios explicitly requires a small learning rate *α*.

### Simulations of learning

The previous sections revealed what to expect from training the learning rules 2 and 3 on certain types of reward. Specifically, we investigated the connection strengths *G* and *N* after many experiences of either totally predictable or totally random rewards. In this section, we aim to confirm and extend those results using numerical simulations rather then analytic methods.

Fig 5 shows the results of simulating the gradual change of connection weights in four different tasks. In all those simulations, *G* and *N* change according to the learning rules 2 and 3. The parameters we used roughly fulfill the conditions 12 and 13 for learning of the correct magnitudes of payoffs and costs^4^.

**Fig 5.**
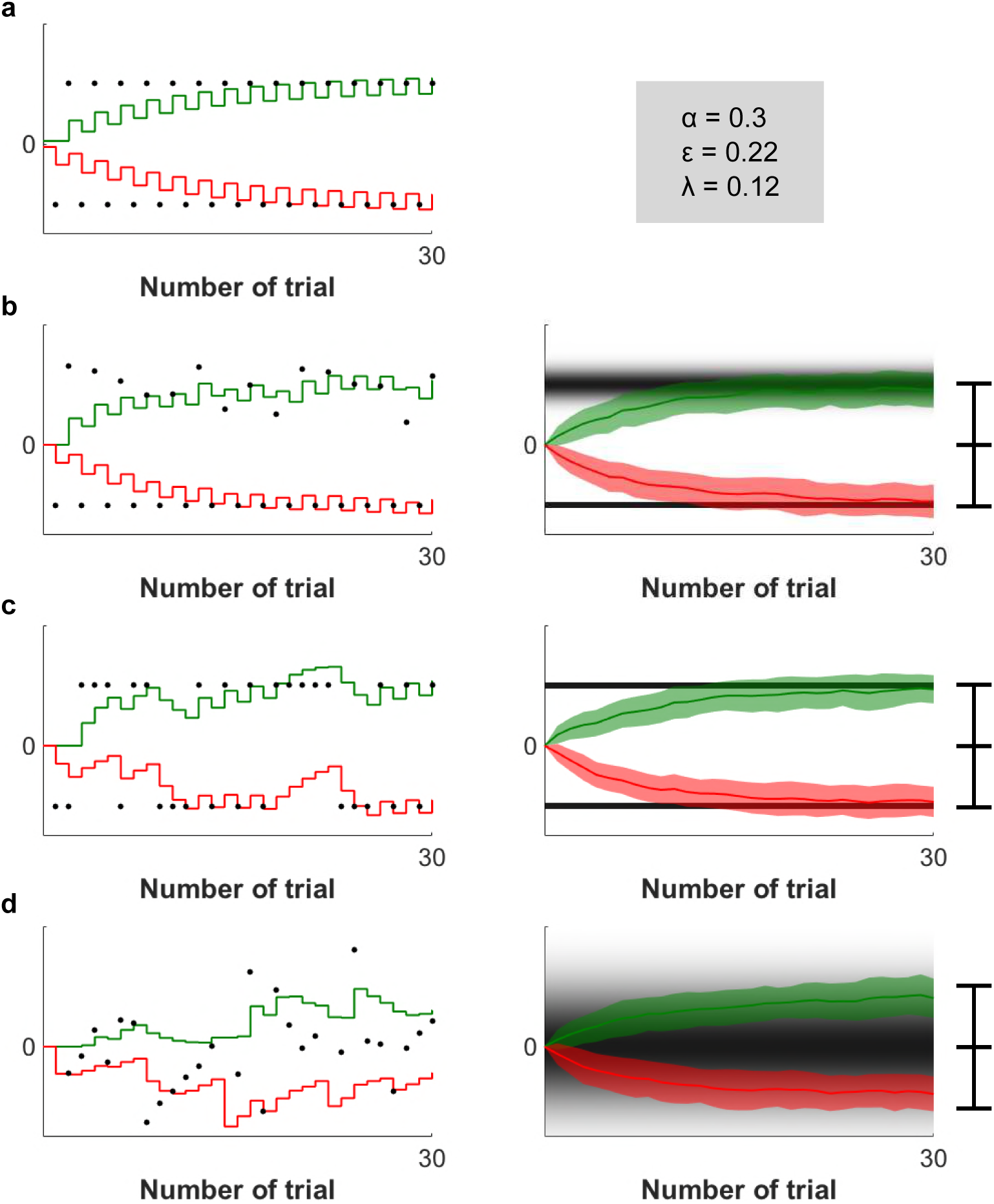
Simulations of learning. In all graphs, the collective strength *G* of the Go weights is depicted in green, while the negative collective strength −*N* of the No-Go weights is depicted in red. The received rewards are indicated by black dots in the panels on the left, while the underlying reward distributions are represented by gray background shadings in the panels on the right. Each simulation shows how *G* and *N* change due to the reception of 30 prediction errors. Panel (a) contains a simulation based on predictable, alternating rewards. It also contains the parameter values used for the simulations. Panels (b) to (d) show both single and averaged simulations of stochastic rewards. The shaded areas around the averages of *G* and *N* in the right column indicate one standard deviation. The bars behind the averaged simulations indicate the mean and mean spreads of the respective reward distributions.

The simulation in Fig 5a is based on a repeating an action that reliably results in a cost −*n*, followed by a payoff *p*. An analytic treatment of that case can be found in the previous sections. Both weights constantly oscillate due to the alternation of payoff and costs. This oscillating behavior is superimposed with learning curves that take the weights from their initial values towards the magnitudes of the payoffs and costs respectively. After 30 trials, *G* and *N* represent good approximations of *p* and *n*.

Fig 5b is similar to Fig 5a, with a slight variation: Just as in Fig 5a, payoffs and costs alternate reliably. But while the cost is again held constant at −*n*, this time the payoff *P* is sampled from a fixed distribution in each trial. Thus, the task includes both stochastic and deterministic components: each repetition of an action results in a fixed cost, which is followed by an uncertain reward. The depicted simulations show that under such conditions, *N* eventually represents the cost *n*, while *G* converges towards the mean payoff 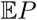.

Finally, panels 5c and 5d contain simulations of repeated actions with rewards drawn at random from fixed distributions. We simulate the experience resulting from such actions by sampling rewards from a fixed distribution on each trial. The stochastic nature of this procedure causes the evolution of the weights *G* and *N* to be different each time the simulation is run. To overcome that effect and segregate random fluctuations from reproducible effects, we collect and average a large number of runs. Each row in Fig 5b - d contains both a single run of the simulation and an average over 500 successive runs. In the above sections, we proved that in purely stochastic tasks, the weights would approximate key statistics of the reward distribution after convergence. Those statistics are indeed approximated in the simulations, confirming the results of the analytic treatment above.

### Simulations of the effect of dopamine depletion

In the previous sections, we focused on the change of the synaptic weights associated with a single action during the accumulation of experience. In this section, we redirect our attention. Instead of considering one action during learning, we now consider multiple actions after learning, and ask: can effects of dopamine depletion on choice behavior be explained in terms of payoffs versus costs?

In a classic experiment illustrated in Fig 6a, rats were given a choice between pressing a lever in order to obtain a nutricious pellet, and freely available lab chow [23]. Normal animals were willing to work for pellets, but after dopamine depletion they were not any more willing to make an effort and preferred a less valuable but free option. Collins and Frank [8] provided a mechanical explanation for this surprising effect. The theory proposed in this paper accounts for it in a conceptually similar but slightly simpler way. Here, we explain our modeling of the experiment, and then describe the simulations - the differences to the account of OpAL model are presented in Discussion.

**Fig 6.**
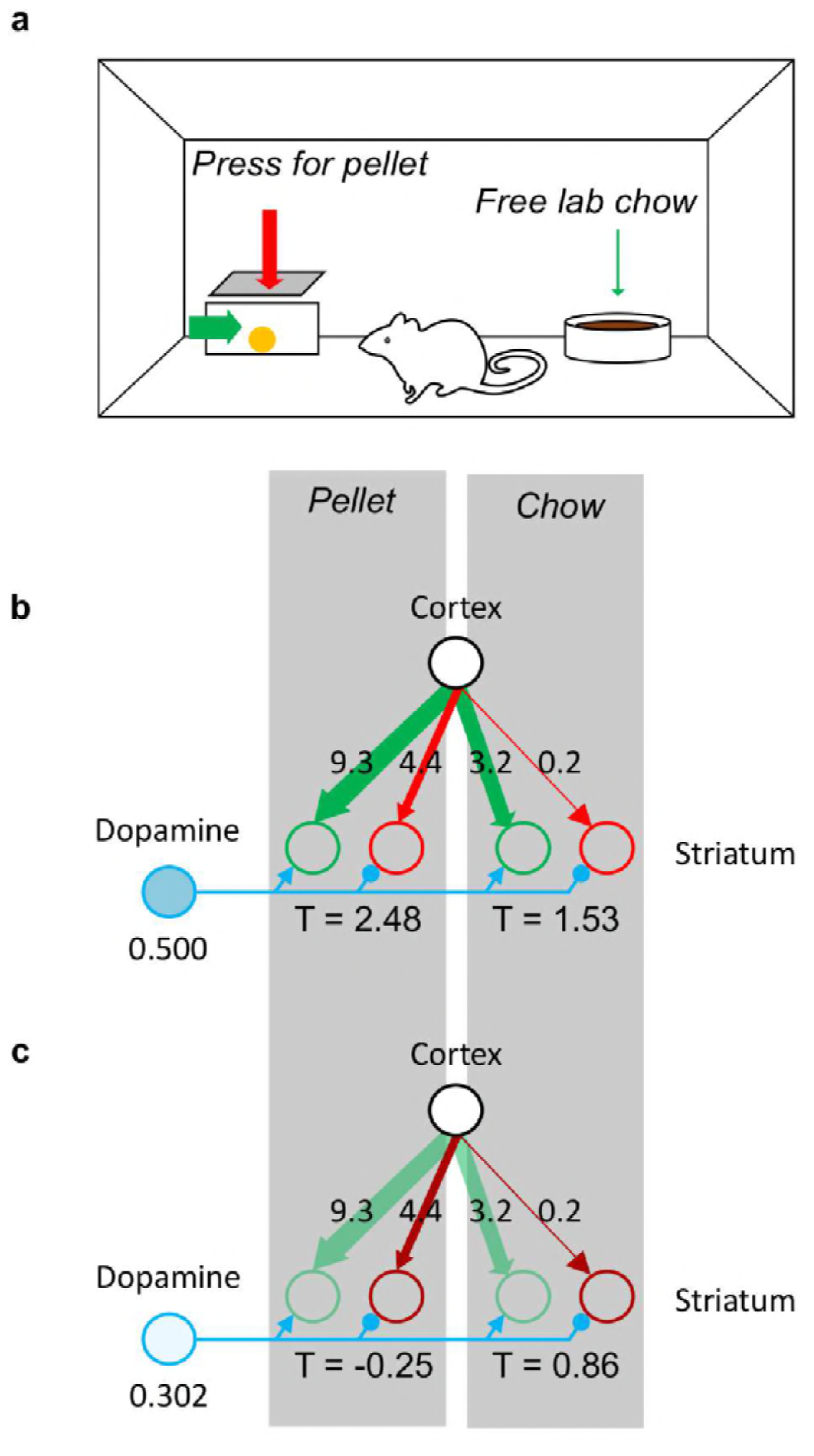
Effects of dopamine depletion on the willingness to exert effort. (a) Schematic illustration of the experimental setup. (b) Action selection in dopamine intact state. Green and red circles on the left denote striatal Go and No-Go neurons associated with pressing the lever, while the green and red circles on the right denote the neurons associated with approaching free food. The strength of the synaptic connections, which result from simulated learning, are indicated by the thickness of the arrows, and labels. The parameters used for the simulations were obtained through a fit of the model to the experimental data. The blue circle represents a population of dopaminergic neurons, and its shading indicates the level of activity. (c) Action selection in dopamine depleted state. The notation is the same as in panel B, but additionally the light green color of the connection of the Go neurons indicates that their gain has been reduced, while the dark red color of the connections of the No-Go neurons symbolizes an increased gain.

To model the experiment, we need to specify how the striatal weights *G* and *N* and the motivation signal affect the output of the basal ganglia system, and how that output then affects choice. We refer to the output of the basal ganglia as the thalamic activity, denoted by *T*. *T* depends on the cortico-striatal weights *G* and *N*, and dopaminergic motivation signal denoted by *D*. Even though this relationship might admittedly be complex, we restrict ourselves to just capture the signs of the dependencies by using a linear approximation:

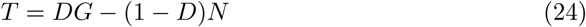

In the above equation, the first term *DG* corresponds to input from the striatal Go neurons. This term is positive, because the projection from striatal Go neurons to the thalamus involves double inhibitory connections (see Fig 1) resulting in an overall excitatory effect. The activity of the Go neurons depends on synaptic weights *G*. We assume that their gain is modulated by the dopaminergic input *D*, based on the observation of an increased slope of the firing-input relationship in the presence of dopamine [24]. The second term −(1 − *D*)*N* corresponds to input from the striatal No-Go neurons. It has a negative sign because the projection form the No-Go neurons to the thalamus includes three inhibitory connections. The activity of the striatal No-Go neurons depends on their synaptic weights *N*, and we assume that their gain is reduced by dopamine, so the synaptic input is scaled by (1 − *D*). In Eq (24), we assume that *D* ∊ [0, 1], and that the value of *D* = 0.5 corresponds to a baseline level of dopamine for which both striatal populations equally affect the thalamic activity. Although arising from a slightly different induction, the action value defined by Eq 24 is directly proportional to the action value proposed by Collins and Frank, which is defined by Eq 4 of their publication [8]: *Q* ∝ *β_G_G* − *β_N_ N*. One easily verifies the direct proportionality of the two expressions by rewriting *D* = 1/2 (1 + (*β_G_* − *β_N_*) / (*β_G_* − *β_N_*)).

How does thalamic activity affect choice? Again, we use a very simple dependency to capture the key aspects of that relationship: In our model of the experiment, we calculate the thalamic activity for each option. Then, we add some random noise independently to each option. Finally, all options with negative noisy thalamic activity are discarded, and the option with the highest noisy thalamic activity is chosen. If the noisy thalamic activity is negative for all available options, no choice will be made; the model defaults to staying inactive.

Often in similar situations, the softmax rule is the preferred choice procedure. According to that rule, one should first transform the set of different action values (or thalamic activities in this case) into a probability distribution over the available actions, by use of the softmax function. Then, one should sample an action from that distribution, and declare it the choice of that trial. Collins and Frank’s OpAL model [8] exemplifies the use of the softmax rule.

We deliberately decided against this conventional approach and in favor of the above described procedure to accommodate a certain feature of the data presented in [23]: The dopamine depleted group of rats differed from the control group not only in their willingness to work for food, but also in their overall food consumption. The dopamine depleted rats consumed less food in total (see Fig 7c). We can hope to capture this effect with our model, since it allows for the possibility to make no choice at all, and thus consume neither of the food items. A softmax decision rule, on the other hand, forces a choice on each trial, and must therefore always lead to the same number of consumed food items.

**Fig 7.**
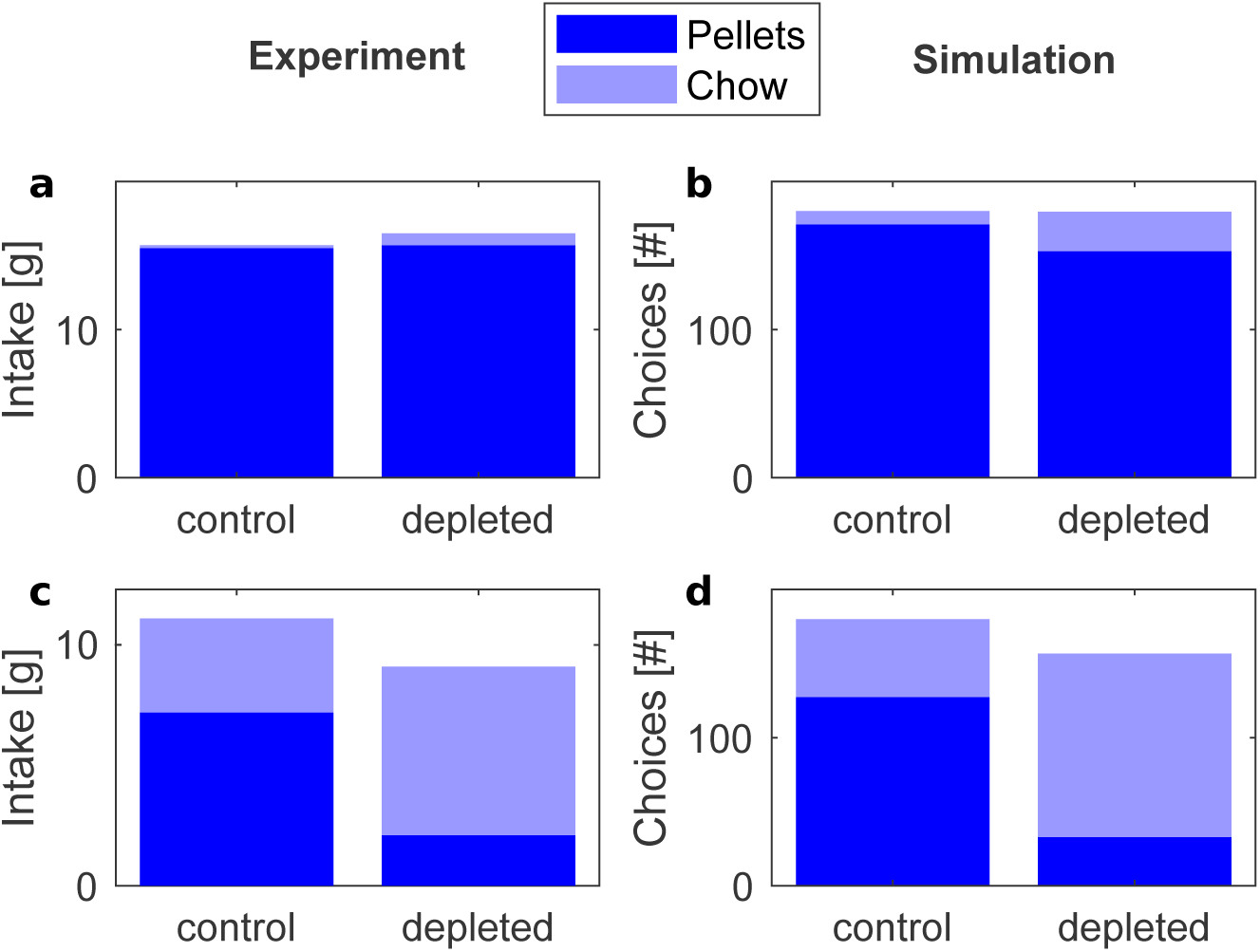
Frequency of choosing pellets and lab chow in dopamine intact (dark blue) and dopamine depleted (light blue) states. The top displays (a) and (b) correspond to a condition with free pellets, while the bottom displays (c) and (d) correspond to a condition where pressing a lever was required to obtain a pellet. The left displays (a) and (c) re-plot experimental data. The values in those displays were taken from Figures 1 and 4 respectively in the paper by Salamone et al. The right displays (b) and (d) show the results of simulations. The parameters used to simulate learning were *α* = 0.1, *ϵ* = 0.6327 and *λ* = 0.0204.

Fig 6b illustrates how the model can account for the behaviour when the dopamine level has a normal baseline value. In the figure, the strength of the cortico-striatal connections is denoted by the labels and the thickness of arrows. Pressing the lever gives a high payoff, so the weights of Go neurons selective for this action are strong, but it also has a substantial cost, so the No-Go weights are also present. On the other hand, the free food is not particularly nutritious so the Go weights are weak, and there is no cost, so the No-Go weight is negligible. When the dopamine level is at baseline, the positive and negative consequences are weighted equally, so the thalamic neurons selective for pressing the lever have overall higher activity, which ultimately leads to a high likelihood for this action to be chosen over the free option. By contrast, Fig 6c shows that when the dopamine level is reduced, costs are weighted more than payoffs, and the thalamic activity associated with pressing the lever drastically decreases. Approaching free food has only negligible cost; therefore, the activity of thalamic neurons selective for this option is now higher, and this action is overall more likely to be chosen.

A quantitative fit of our model to Salamone et al.’s experimental results [23] is illustrated in Fig 7. The panels on the left side in Fig 7 summarize experimental data: the top-left display corresponds to a condition in which both high-valued pellets and the low-valued lab chow were freely available. In this case, the animals preferred pellets irrespectively from dopamine level. The bottom-left panel corresponds to the condition in which the animal had to press a lever in order to obtain a pellet, and as mentioned before, after injections of a dopamine antagonist they started to prefer the lab chow.

In our model of the experiment, we run through a sequence of trials mimicking those illustrated in Fig 6: on each trial, the model makes a choice between two actions - pressing a lever or approaching lab chow - or remains inactive. Before the main experiments, the animals were trained to press lever to obtain reward and were exposed to the lab chow [23]. To parallel this in simulations, the model was first trained such that it experienced each action a number of times, received corresponding payoffs and costs, and updated its weights according to equations 2 and 3. The weights resulting from that learning are reported in Fig 6b and Fig 6c. Then, the model was tested with baseline and reduced dopaminergic motivation signal. As described in Materials and Methods, the parameters of the model were optimized to match experimentally observed behavior. As shown in the right displays in Fig 7, the model was able to reproduce the observed pattern of behavior. This illustrates model’s ability to capture both learning about payoffs and costs associated with individual actions and the effects of the dopamine level on choices.

### An actor-critic variation

So far, we assumed that the reward prediction is computed by the same striatal neurons that encode the payoffs and costs of actions. Only one network was involved: that which is responsible for the choice of action. We refer to such a network as ‘actor’ in the remainder of this exposition. In this section, we look at how the theory described above generalizes to the actor-critic framework [26]. That framework assumes that the reward prediction is not computed by the actor, but by a separate group of striatal patch neurons called the ‘critic’. More formally, the purpose of that critic is to learn the value *V* of the current state.

One way to generalize our theory in this direction is to keep the actor network unaltered, while substituting it with a similar critic network that learns by the very same rules 2 and 3:

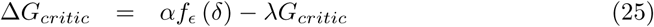

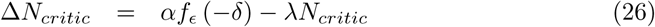

The crucial difference between the actor and the critic is that the critic network is not selective for the action, but only for the state. It thus learns the value of a state irrespective of the actions chosen. Importantly, the critic is in charge of suppling the reward predictions. Those predictions are compared to the actual outcomes to produce the reward prediction errors *δ* from which both networks learn.

We take the state value to be encoded in the difference of *G_critic_* and *N_critic_*: *V_critic_* = 1/2(*G_critic_* − *N_critic_*). The change of the state value on each trial can be obtained by subtracting equations 25 and 26:

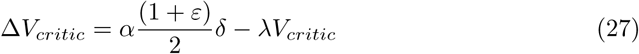

The prediction error *δ* - which teaches the actor as well - is the difference between the obtained reward *r* and the reward prediction by the critic:

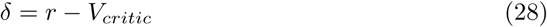

What would be learned with that architecture? If the same action is selected on each trial, the actor will learn in exactly the same way as the critic. Then, the prediction error in the actor-critic model is the same as in the actor-only model described above, and the weights of the actor in the actor-critic model converge to exactly the same values as for the actor-only model. However, this reasoning does not seem to apply if more than one action is available: empirically, animals then select the actions that maximize their rewards in their own perception. In the process of learning, they will likely sample all available actions.

If such behavior generates input for an actor-critic model, the critic will integrate the experience of all those trials, and will thus represent a mixture of the expected rewards associated with the available actions. This generally interferes with correct learning of the payoffs and costs of the different actions. However, there is a caveat: one of the available actions will eventually proof most useful; as soon as the animal has determined that best action, it will select it in the majority of cases. That, in turn, forces the critic into mainly representing the expected reward of this best action. As a final consequence, also payoff and cost of that best action are inferred correctly.

We confirmed the conclusions of this discussion empirically for the model specified above: in Fig 8, we present simulations of a task in which the subject must choose between two actions. Both actions reliably yield a constant cost followed by a constant payoff each time they are selected. One of the actions is unambiguously superior to the other: its payoff is larger and the its cost is lower.

**Fig 8.**
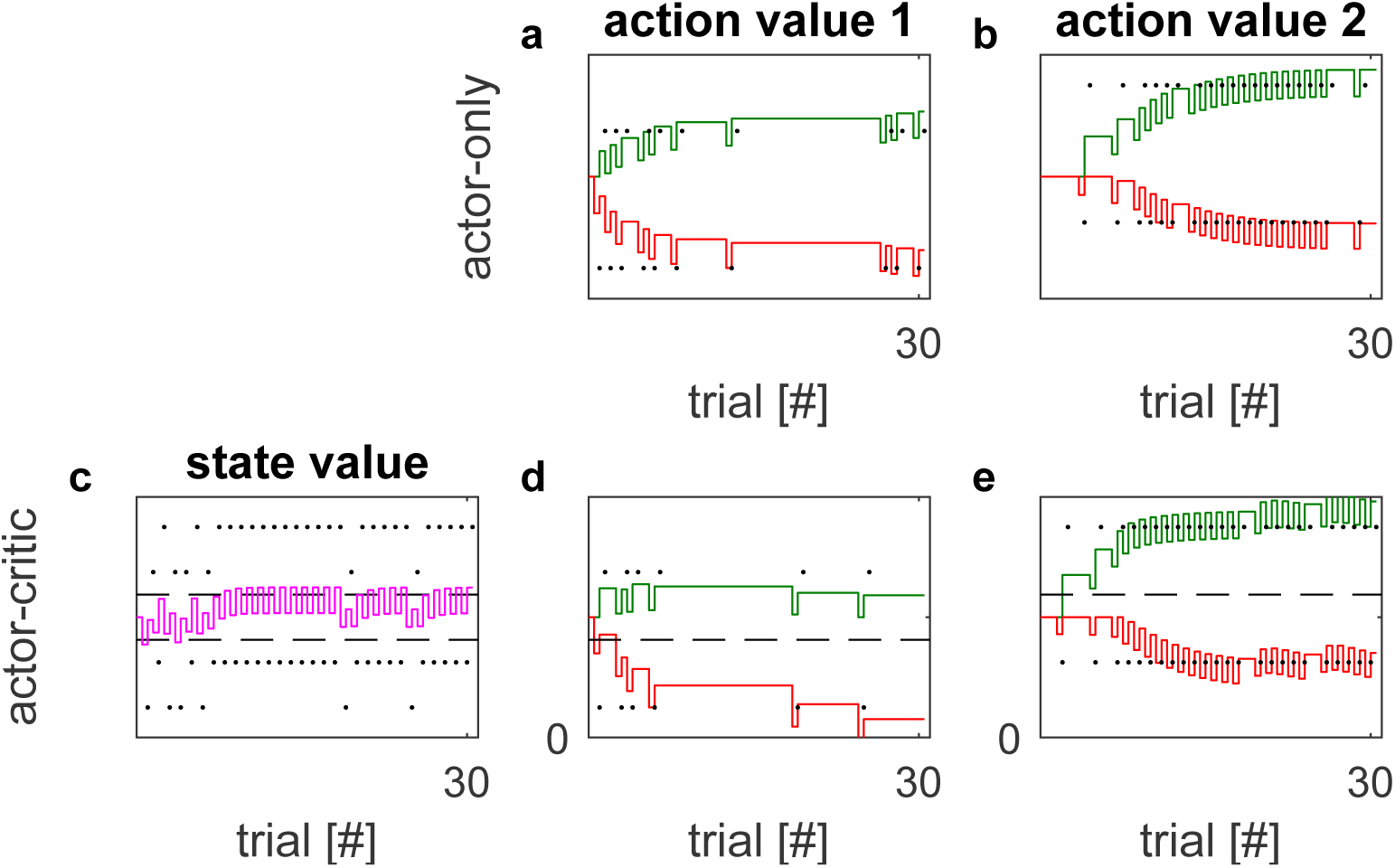
Actor-only in comparison with actor-critic learning. The columns labeled with ‘action value 1’ and ‘action value 2’ show the simulated evolution of the collective synaptic weights *G* and *N* of the actor network over 30 successive trials. The weights *G* are drawn as solid green lines, the negative weights *N* are drawn as solid red lines. The rewards obtained by choosing the respective actions are indicated by black dots. For the actor-critic simulations (second row), we additionally provide the evolution of the state value in panel c. There, the state value *V_critic_* is represented by a solid purple line. The expected rewards of both actions are indicated by dashed horizontal lines. The parameter settings used in these simulations were *α* = 0.4, *ϵ* = 0.519, *λ* = 0.1013 and *β* = 0.9. The same set of parameters was used for both the actor-only and the actor-critic model.

Both an actor-only model and an actor-critic model interacted with that task. On each trial, an action was selected by sampling from a softmax distribution over all available actions: the probability of choosing action *a* was proportional to exp (*βQ_a_*), where *Q_a_* = 1/2 (*G_a_* − *N_a_*) was the action value, and *β* was the softmax temperature. Fig 8 shows the temporal evolution of the involved synaptic weights over the course of learning. Panels 8a and 8b depict the actor-only evolution of the weights *G* and *N* that encode the payoffs and costs of of actions 1 and 2, respectively. For both actions, payoffs and costs are learned correctly. Learning is notably slower for action 1. This is easily explained: action 1 is the worse of the two options, and thus chosen much less frequent. In contrast, the actor-critic driven evolution of the same weights presented in panels 8d and 8e leads to a correct estimate of the payoff and cost only for the superior action 1. Learning is impaired for the inferior action 2, as anticipated in the qualitative discussion above. The state value, presented in panel 8c, provides further confidence in the validity of that discussion: Instead of encoding a mixture of the values of all available actions, it converges to the value of the superior action, indicated by the higher of the two dashed lines.

## Discussion

This article describes how the positive and negative consequences of actions can be separately learned on the basis of a single teaching signal encoding reward prediction error. In this section we relate the theory with data and other models, state experimental predictions, and highlight the directions in which the theory needs to be developed further.

### Relationship to experimental data

The model described in this paper was shown in simulations to avoid action requiring effort when the motivational signal was reduced. The unwillingness to make an effort for reward in dopamine depleted state has also been observed in other paradigms: During a choice in a T-maze, dopamine depleted animals were less likely to go to an arm with more pellets behind the barrier, but rather chose the arm with easily accessible but fewer pellets [27]. Parkinson’s patients were not willing to exert as much physical effort by squeezing a handle in order to obtain reward as healthy controls, especially if they were off medications [28]. These effects can be explained in an analogous way [8] by assuming that in the dopamine depleted state the effort of crossing the barrier or squeezing a handle is weighted more, resulting in lower activity of thalamic neurons selective for this option. Both in OpAL and the model proposed here, reducing the dopamine level reduces the tendency to choose actions involving costs, and thus changes preferences.

Let us now consider how the weight changes in our model relate to known data on synaptic plasticity in the striatum. Fig 9b illustrates the weight changes when an animal performs an action involving a cost *n* in order to obtain a payoff *p* (Fig 9a), e.g. pressing a lever in order to obtain a pellet. The direction of changes in *G* and *N* depending on the sign of *δ* are consistent with the changes of synaptic weights of Go and No-Go neurons observed at different dopamine concentrations. Fig 9c shows experimentally observed changes in synaptic strengths when the level of dopamine is low (displays with white background) and in the presence of agonists (blue background) [11]. Note that the directions of change match those in the corresponding displays above, in Fig 9b.

**Fig 9.**
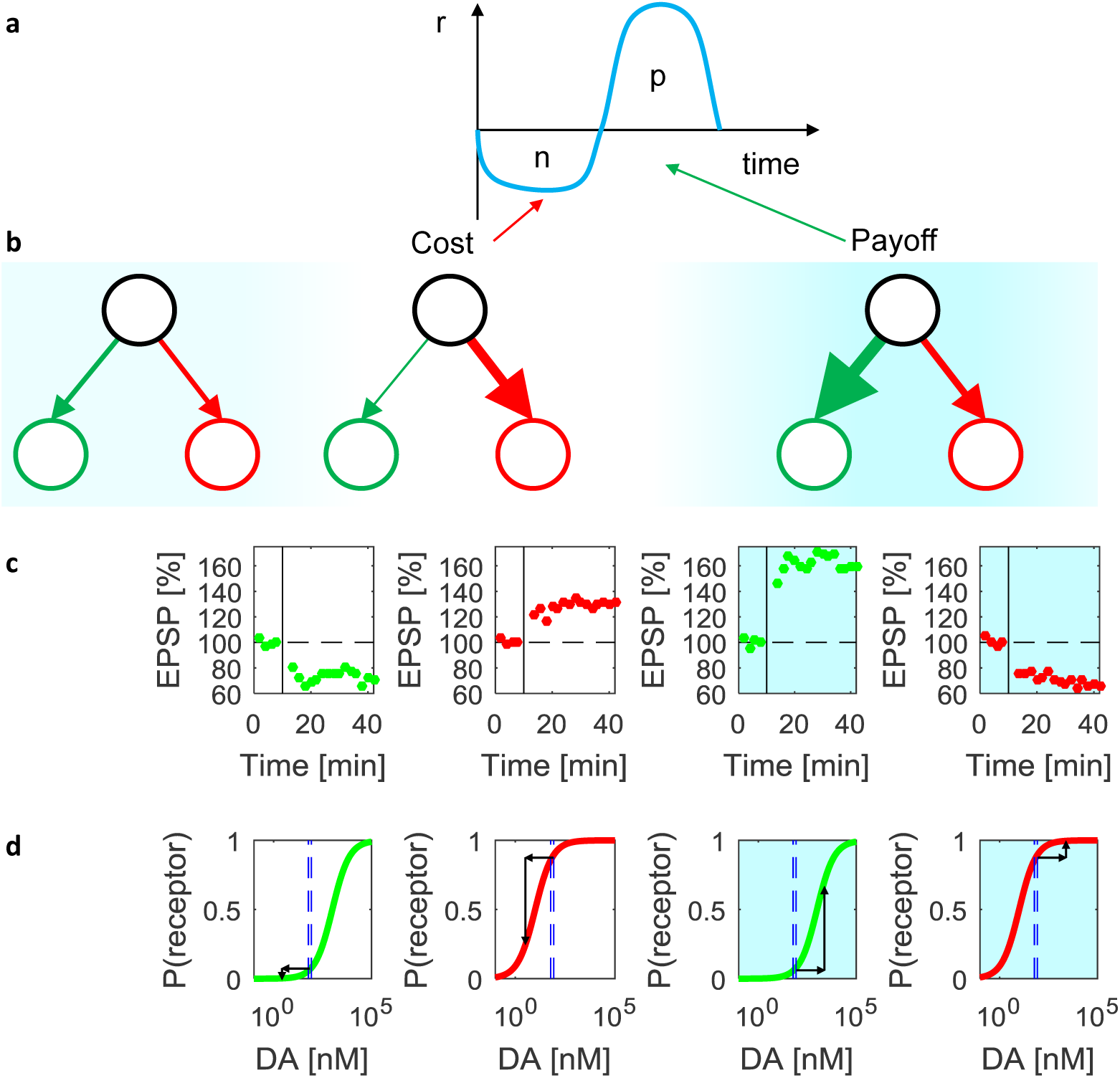
Relationship of learning rules to synaptic plasticity and receptor properties. (a) Instantaneous reinforcement *r* when an action with effort *n* is selected to obtain payoff *p*. (b) Cortico-striatal weights before the action, after performing the action, and after obtaining the payoff. Red and green circles correspond to striatal Go and No-Go neurons, and the thickness of the lines indicates the strength of synaptic connections. The intensity of the blue background indicates the dopaminergic teaching signal at different moments of time. (c) The average excitatory post-synaptic potential (EPSP) in striatal neurons produced by cortical stimulation as a function of time in the experiment by [11]. The vertical black lines indicate the time when the synaptic plasticity was induced by successive stimulation of cortical and striatal neurons. The amplitude of EPSPs is normalized to the baseline before the stimulation indicated by horizontal dashed lines. The green and red dots indicate the EPSPs of Go and No-Go neurons respectively. Displays with white background show the data from experiments with rat models of Parkinson’s disease, while the displays with blue background show the data from experiments in the presence of corresponding dopamine receptor agonists. The four displays re-plot the data from Figures 3E, 3B, 3F and 1H in the paper by [11]. (d) Changes in dopamine receptor occupancy. The green and red curves show the probabilities of D1 and D2 receptor occupancies in a biophysical model of [29]. The two dashed blue lines in each panel indicate the levels of dopamine in dorsal (60 nM) and ventral (85 nM) striatum estimated on the basis of spontaneous firing of dopaminergic neurons using the biophysical model [30]. Displays with white and blue backgrounds illustrate changes in receptor occupancy when the level of dopamine is reduced or increased respectively.

These directions of changes in striatal weights are also consistent with other models of the basal ganglia [8, 12], but the unique prediction of the rules described in this paper is that the increase in dopaminergic teaching signal should mainly affect changes in *G*, while the decrease in dopamine should primarily affect *N*. Thus, the dopamine receptors on the Go and No-Go neurons should be most sensitive to increases and decreases in dopamine level respectively. This matches with the properties of these receptors. The D2 receptors on No-Go neurons have a higher affinity and therefore are sensitive to low levels of dopamine compared to D1 receptors on Go neurons [31]. This property is illustrated in Fig 9d where the green and red curves show the probabilities of D1 and D2 receptors being occupied as a function of dopamine concentration. The blue dashed lines indicate the levels of dopamine in the striatum predicted to result from spontaneous firing of dopaminergic neurons [30]. At these levels most D1 receptors are deactivated. Thus the D1 receptor activation will change when the dopamine goes up, but not when it goes down, as indicated by the black arrows. This is consistent with the stronger impact of positive prediction errors on the weight changes of the Go neurons implemented in Equation 2. By contrast, the D2 receptors are activated at baseline dopamine levels, so their activation is affected by the decreases in dopamine level but little by increases, in agreement with stronger impact of positive prediction errors on the No-Go neurons implemented in Equation 3. In summary, the plasticity rules allowing learning positive and negative consequences are consistent with the observed plasticity and the receptor properties.

Recently, there has been a debate concerning the fundamental concept of basal ganglia function, i.e. the relationship between the Go and No-Go neurons: on one hand they have the opposite effects on a tendency to make movements [2], but on the other hand they are co-activated during action selection [33]. The presented theory is consistent with both observations: It assumes that Go and No-Go neurons have opposite effects on movement initiation. But during action selection the basal ganglia need to calculate the utility which combines information encoded by both populations, so may require their co-activation.

The proposed model assumes that while an animal makes an effort, the reward prediction error should be negative, thus the dopamine level should decrease. However, at the time of lever pressing the system needs to be energized to perform a movement, so one could expect increased level of dopamine. Furthermore, voltametry studies measuring dopamine concentration in striatum did not observe decrease in dopamine level during lever pressing [32]. Nevertheless a recent study recording activity of single dopaminergic neurons that provided a better temporal resolution reported that dopaminergic neurons increased the activity before movement, and then decreased it below baseline during movement [30]. The increase before movement may be related with energizing system for movement, while the decrease during movement may be related with representing effort.

Another study [34] directly tested whether dopaminergic signals encode expected efforts alongside expected payoffs. It reports dopaminergic bursts in the nucleus accumbens of rats, triggered by unexpected opportunities. According to the theory of temporal difference learning, such bursts encode reward prediction errors. These prediction errors occur whenever a reward or the anticipation of a reward is unexpectedly encountered. If, for instance, an unexpected cue signals an opportunity to gain reward, a prediction error equal to the value of the opportunity will arise. In our theory, the value of opportunities or actions is assembled from payoffs, costs and motivation. Does the dopaminergic signal investigated in [34] signal the value of opportunities according to our theory? Three different opportunities featured in the investigation of Hollon et al.: an opportunity that yielded a small payoff for little effort served as a reference, and was compared with two high-payoff opportunities. Those options required different levels of effort, chosen such that the rats would prefer one of them over the reference, while rejecting the other one. In economic terms, one opportunity had a higher utility than the reference option, while the other one had a lower utility. The dopamine measurements obtained by Hollon et al. did not reflect the value of actions - they were much better explained by the mere payoff of the opportunities. Dopamine concentrations were uniformly higher for the high-payoff opportunities than for the reference option, even though their values (expressed through the choices of the rats) were spread around the value of the reference option. The different efforts showed strongly in the choice, but only negligibly in the dopamine bursts. This poses a challenge to our theory, which assumes that both payoffs and costs of actions are encoded in the basal ganglia. However, from the results of the study by Hollon et al. it is not clear if effort is not represented in the basal ganglia at all, or simply does not affect the value signalled by dopaminergic neurons. Distinction between these hypotheses will require further experiments, as discussed below.

### Experimental predictions

A direct test of the proposed model could involve recording of activity of Go and No-Go neurons (e.g. with photometry) during a task in which an animal learns the payoffs and costs associated with an action. Assuming that *G* and *N* are reflected in the activity of the Go and No-Go neurons while the animal evaluates an action (i.e. just before its selection), one could analyze the changes in the activity of Go and No-Go neurons across trials. One could compare if they follow the pattern predicted by the rules given in this paper, or rather by other rules proposed to describe learning in striatal neurons [7, 8, 14].

Similarly as the OpAL model [8], the theory proposes that the positive and negative consequences are separately encoded by the Go and No-Go neurons which are differentially modulated by dopamine. The theory predicts that agonists specific to just one of the striatal populations (e.g. a D2 agonist), should decrease the effect of consequences encoded by this population (e.g. negative) without changing the impact of the other population. This prediction could be tested in an experiment involving choice between options with both payoff and cost. In particular, the theory predicts that the degree of preference of a neutral option (*p* = 1*, n* = 1) over a high cost option (*p* = 1*, n* = 2) should increase with D2-agonist, while the preference of a high payoff option (*p* = 2*, n* = 1) over a neutral option (*p* = 1*, n* = 1) should not be affected by the D2-agonist.

It could also be worthwhile to investigate whether changing the influence of positive and negative consequences on choice can not only be achieved by pharmacological manipulations, but also by changing a behavioral context such as hunger, or reward rate which has been shown to affect the average dopamine level [19]. If such an experiment was done in humans (or non-human primates), an eye-tracker could be used to investigate whether participants spend more time on a part of the stimulus informing about payoff in blocks with high hunger or reward rate.

The theory assumes that the synaptic plasticity rules include a decay term proportional to the value of the synaptic weights themselves. Decay terms are also present in other models of learning in basal ganglia [15, 35, 37]. This class of models predicts that the synaptic weights of striatal neurons which are already high increase less during potentiation than the smaller weights (an opposite prediction is made by the OpAL model [8], where the weights scale the prediction error in the update rule). This prediction could be tested by observing the Excitatory Post-Synaptic Currents (EPSCs) evoked at individual spines. The class of model including decay predicts that the spines with smaller evoked EPSCs before inducing plasticity should be more likely to potentiate.

### Relationship to other theories

The proposed model builds on the seminal work of Collins and Frank [8], who proposed that the Go and No-Go neurons learn the tendency to execute and inhibit movements, and how the level of dopamine changes the influence of the Go and No-Go pathways on choice. The key new feature of the present model is the ability to learn both payoffs and costs associated with a single action. We demonstrated above that when the model repeatedly selects an action resulting first in a cost and then in the payoff, *G* and *N* - under certain conditions that we specified - converge to the magnitudes of that payoff and cost. This is not so in the original OpAL model, as we shall show in a brief analysis.

Collins and Frank [8] demonstrated that when the environment is stationary and prediction error *δ* converges to zero, then the weights *G* and *N* in the OpAL model converge to bounded values. However, we will show that Go and No-Go weights converge to zero when an action that results first in a cost and then in the payoff is repeatedly selected.

The OpAL model is based on the actor critic framework; hence, the prediction error is defined as in Eq (28). The weights of the critic are modified simply as Δ*V* = *αδ*. The weights of the actor are modified according to the following equations [8]:

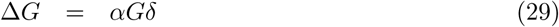

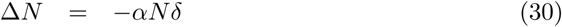

Fig 10 shows how the weights change in a simulation of the OpAL model. The weights of the critic approach a value close to the average of payoff and cost. Let us consider what happens in the model once the critic weight stops changing between trials (i.e. from ~ 10th trial onward in Fig 10). The weight of the critic still changes within a trial, i.e. decreases when cost is incurred and increases after a payoff. This happens because the prediction error oscillates around 0, i.e. it is equal to *δ* = −*d* while incurring a cost and *δ* = *d* while receiving a payoff, where *d* is a constant. If so, let us consider how a Go weight changes within a trial. According to Eq (29) the weight changes as follows:

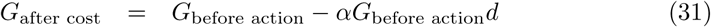

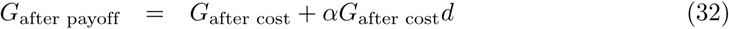

**Fig 10.**
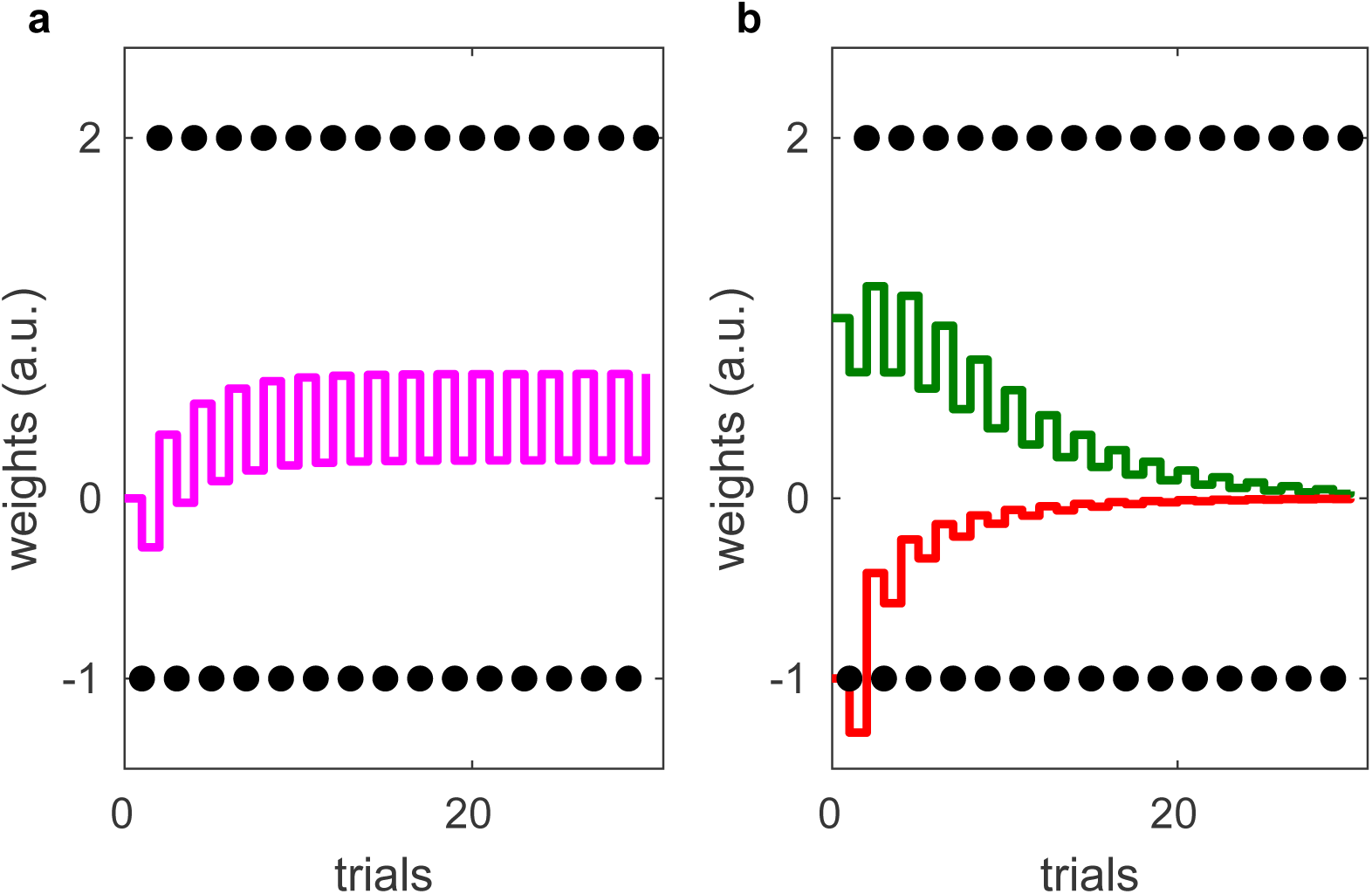
Changes in the weight *G* of Go neurons, *N* of No-Go neurons and *V* of the critic in the OpAL model over the course of simulations. (a) The purple line represents the evolving critic weight. The experienced rewards are indicated by black dots. (b) The actor weights, represented by a green and a red line respectively, were initialized to *G* = *N* = 1. Again, the black dots indicate the received rewards. The simulation was run with learning rate *α* = 0.3.

Substituting Eq (31) into Eq (32) we obtain:

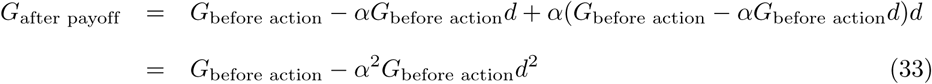

We see that within a trial a Go weight decays proportionally to is value, resulting in an exponential decay across trials seen in Fig 10. Analogous calculations show that the No-Go weight decays in the same way. We conclude that the OpAL model is unable to estimate positive and negative consequences for actions which result in both payoffs and costs. It is worth noting that the decay of actor weights to zero demonstrated above is specific to the version of basal ganglia model proposed by Collins and Frank [8], but would not be present in another version of the model [35] where the learning rules include a special term preventing the weights from approaching zero.

The model described in this paper has been shown to account for the effects of dopamine depletion on willingness to make effort, which have also been simulated with the OpAL model. To simulate the effects of dopamine depletion on choice between an arm of a T-maze with more pellets behind a barrier and an arm with with fewer pellets, [8] trained a model on three separate actions: eating in the left arm, eating in the right arm, and crossing a barrier. In this way it was ensured that each action had just payoff or just cost, and the model could learn them. Subsequently, during choice the model was deciding between a combination of two actions (e.g. crossing a barrier and eating in the left arm) and the other action. By contrast, the model proposed in this paper was choosing just between the two options available to an animal in an analogous task (Fig 6), because it was able to learn both payoffs and costs associated with each option. This is a useful ability, as most real world actions have both payoffs and costs.

In the original paper introducing the plasticity rules [16], it was proposed that the rules allow the Go and No-Go neurons to encode reward variability, because when an action results in variable rewards, both *G* and *N* increase during learning. It was further proposed that the tonic level of dopamine controls the tendency to make risky choices, as observed in experiments [36], because it leads to emphasizing potential gains, and under-weighting potential losses. However, here it is proposed that the striatal learning rules primarily sub-serve a function more fundamental for survival, i.e. learning payoffs and costs of actions. From this perspective, the influence of dopamine level on tendency to make risky choices arises as a by-product of a system primarily optimized to weight payoffs and costs according to the current motivational state.

### Directions for the future work

There are multiple directions in which the presented theory could be extended. For example, the theory has to be integrated with the models of action selection in the basal ganglia to describe how the circuit selects the action with the best trade-off of payoffs and costs. Furthermore, the theory may be extended to describe the dependence of the dopaminergic teaching signal on the motivational state [38].

It is intriguing to ask whether the evaluation of actions combining separately encoded positive and negative consequences is also performed by areas beyond the basal ganglia. Indeed, positive and negative associations are encoded by different populations of neurons in the amygdala [39]. Moreover, an imaging study [40] suggests that costs and payoffs are predicted by the amygdala and the ventral striatum respectively, and ultimately compared in the prefrontal cortex. Furthermore, different cortical regions preferentially project to Go or No-Go neurons [41], raising the possibility that the positive and negative consequences are also encoded separately in the cortex. Therefore, it seems promising to investigate if similar plasticity rules could also describe learning beyond the basal ganglia.

## Materials and methods

During simulations of an experiment by Salamone et al. [23], the model received payoff *p*_pellet_ = 10 for choosing a pellet, and payoff *p*_chow_ for approaching the lab chow. The model was simulated in two conditions differing in the cost of choosing a pellet which was equal to *n*_pellet_ = 0 in the free-pellet condition, and to *n*_pellet_ = *n*_lever_ in a condition requiring lever pressing to obtain a pellet. There was no cost of choosing lab chow (*n*_chow_ = 0). For each condition, the model was simulated in two dopamine states: in the intact state the dopaminergic motivation signal was equal to a baseline value during choice *D* = 0.5 while in the state corresponding to the presence of dopamine antagonist it was set to a lower value *D* = *D_anta_*.

For each condition and state, the behavior of *N_rats_* was simulated. Each simulation consisted of 180 training and 180 testing trials (as each animal in the experiment of [23] was tested for 30 minutes, so 180 trials corresponds to an assumption that a single trial took 10s). At the start of each simulation, the weights were initialized to *G*_pellet_ = *N*_pellet_ = *G*_pellet_ = *N*_pellet_ = 0.1. During each training trial, the model experienced choosing a pellet as well as approaching the lab chow. In detail, it received the cost *n*_pellet_, modified the weights *G*_pellet_ and *N*_pellet_, then received the payoff *p*_pellet_ and modified the weight again, and analogously for the lab chow. During each testing trial, the thalamic activity for each option was calculated from Eq 24), and Gaussian noise with standard deviation *σ* was added. An option with the highest thalamic activity was selected, and if this activity was positive, the action was executed, resulting in the corresponding cost and payoff and weight modification. If thalamic activity for both options was negative, no action was executed and no weights were updated.

The values of model parameters: *p*_chow_*, n*_lever_*, D_anta_, σ* were optimized to match the choices made by the animals. In particular, for each set of parameters, the model was simulated *N_rats_* = 100 times, and the average number of choices 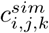 of option *i* in dopamine state *j* and experimental condition *k* was computed. The mismatch with corresponding consumption in experiment 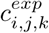 was quantified by a normalized summed squared error:

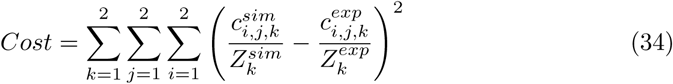

In the above equation 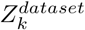 is a normalization term equal to the total number of choices or consumption in a particular condition:

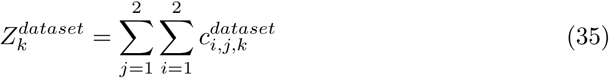

The values of parameters minimizing the cost function were sought using the Simplex optimization algorithm implemented in Matlab, and the following values were found: *p*_chow_ = 3.64, *n*_lever_ = 4.57, *D_anta_* = 0.30 and *σ* = 1.10. Subsequently, the model with these optimized parameters was simulated with *N_rats_* = 6, which was the number of animals tested by [23]. The resulting mean number of choices across animals are shown in Fig 7.

## Acknowledgments

The authors wish to thank Jacqueline Pumphrey for composing a lay summary of this article.

## Footnotes

1 Explicitely, one easily verifies that *f_ϵ_* (*x*) − *f_ϵ_* (−*x*) = (1 + *ϵ*) *x* and *f_ϵ_* (*x*) + *f_ϵ_* (−*x*) = (1 − *ϵ*) |*x*|.

2 Technically, it amounts to *λ/α_Q_* → 0. However, *α_Q_* is an effective learning rate, and so must take values smaller then one. Thus, we really need to let *λ* → 0

3 A small decay is characterized by a decay rate *λ* which is small compared to the learning rate *α*.

4 The parameters are also chosen to facilitate quick convergence. The values presented in Fig 5a mirror that compromise.

